# Assessing antibiotic tolerance of *Staphylococcus aureus* derived directly from patients by the Replica Plating Tolerance Isolation System - REPTIS

**DOI:** 10.1101/2021.05.12.443951

**Authors:** Sebastian C. Herren, Markus Huemer, Claudio T. Acevedo, Federica Andreoni, Alejandro Gómez-Mejia, Srikanth Mairpady Shambat, Barbara Hasse, Reinhard Zbinden, Silvio D. Brugger, Annelies S. Zinkernagel

**Affiliations:** Department of Infectious Diseases and Hospital Epidemiology, University Hospital Zurich, University of Zurich, Switzerland; Institute of Medical Microbiology, University of Zurich, Zurich, Switzerland

## Abstract

Antibiotic tolerant *Staphylococcus aureus* pose a great challenge to clinicians as well as to microbiological laboratories and are one reason for treatment failure. Antibiotic tolerant strains survive transient antibiotic exposure despite being fully susceptible *in vitro*. Thus, fast and reliable methods to detect tolerance in the routine microbiology laboratory are urgently required.

We therefore evaluated the feasibility of the replica plating tolerance isolation system (REPTIS) to detect antibiotic tolerance in *S. aureus* isolates derived directly from patients suffering from different types of infections and investigated possible connections to clinical presentations and patient characteristics.

One hundred twenty-five *S. aureus* isolates were included. Replica plating of the original resistance testing plate was used to assess regrowth in the zones of inhibition, indicating antibiotic tolerance. Bacterial regrowth was assessed after 24 and 48 hours of incubation and an overall regrowth score (ORS) was assigned. Regrowth scores were compared to the clinical presentation.

Bacterial regrowth was high for most antibiotics targeting protein synthesis and relatively low for antibiotics targeting other cellular functions such as DNA-replication, transcription and cell wall synthesis, with the exception of rifampicin. Isolates with a blaZ penicillinase had lower regrowth in penicillin and ampicillin. Low ORSs were more prevalent among isolates recovered from patients with immunosuppression or methicillin-resistant *S. aureus* (MRSA) isolates.

In conclusion, REPTIS is useful to detect antibiotic tolerance in clinical microbiological routine diagnostics. Rapid detection of antibiotic tolerance offers a new diagnostic readout that might allow more tailored treatments in the future.

## Introduction

The human pathobiont and Gram-positive bacterium *Staphylococcus aureus* is recognized as one of the major causes of community associated as well as nosocomial infections (1–3). With the emergence of antibiotic resistance, infections caused by *S. aureus* have become both, a therapeutic challenge and an economic burden for public health. However, antibiotic resistance is not the only cause for treatment failure (4–7). In contrast to the genetically determined antibiotic resistant strains, antibiotic tolerant strains are able to survive antibiotic challenges to which they remain nonetheless fully susceptible. Therefore antibiotic tolerant strains are not able to grow in the presence of antibiotics at therapeutic concentrations (8). While the term antibiotic tolerance encompasses an entire bacterial population, the term persister is used to describe individual bacterial cells that are able to survive antibiotic challenges without being resistant (9).

Strains exhibiting antibiotic tolerance typically go unnoticed in routine diagnostics as they are classified as susceptible. However, recognized as being one reason for treatment failure, many efforts have been undertaken to detect antibiotic tolerance in order to provide more efficient tailored treatments. The minimum bactericidal and inhibitory concentration (MBC/MIC) ratio has been used as a parameter to assess antibiotic tolerance. It describes the discrepancy between the antibiotic concentration required to inhibit growth and the concentration needed to kill 99.9% of the bacterial population. Strains with a MBC/MIC ratio ≥32 are considered antibiotic tolerant (4, 7). However, the MBC/MIC ratio and other assays aiming to detect tolerance such as time-kill curve assays or minimum duration for killing (MDK) measurements (6) are time-consuming and difficult to standardize for every-day routine microbiological diagnostics (10–14).

The need for reliable detection methods of antibiotic tolerance has increased and less laborious alternatives for tolerance detection are being investigated. The tolerance disk test (TD test) as defined by Balaban and colleagues is a modified disk diffusion assay originally designed for *Escherichia coli*. In this assay, antibiotic disks are exchanged with blank disks containing a glucose solution after the formation of an inhibition zone. The addition of glucose facilitates the regrowth of bacteria that survived the antibiotic challenge (persisters) within the inhibition zone (15). Another approach used to detect antibiotic tolerant populations is the replica plating procedure. In a recent study, Matsuo *et al.* introduced the replica plating tolerance isolation system (REPTIS) in order to isolate ciprofloxacin tolerant *S. aureus* in the research laboratory setting (16). Given the complexity of conventional approaches to detect persisters, we aimed to assess the feasibility of this alternative method for tolerance detection translating it from the research laboratory to the routine diagnostic laboratory and to link these findings to clinical parameters. In close cooperation with a diagnostic laboratory, we performed the experiments on bacteria derived directly from patients without any intermediate steps such as freeze-thaw cycles. The results of conventional antibiotic susceptibility testing and REPTIS were compared to each patient’s clinical data.

## Material and Methods

The study was conducted as part of the BacVivo “Bacterial pathogen properties in patient samples” study, a single-center observational study carried out at the University Hospital Zurich, Switzerland. This study had been approved by the cantonal ethics committee, Canton of Zurich, Ethical application number: BASEC 2017-02225. All patients included in this study signed an informed consent.

### Clinical isolates

We screened 126 *S. aureus* strains from clinical samples that were isolated and processed between 2019 and 2020 at the diagnostic laboratory of the Institute of Medical Microbiology (IMM, University of Zurich, Switzerland) in the course of routine diagnostics. All samples were obtained at the University Hospital Zurich (USZ), Switzerland. When detected repeatedly in the same patient, only the first *S. aureus* isolate was included in this study. Out of 126 eligible samples, 125 underwent replica plating. Neither clinical information nor the site of sampling were used as inclusion criteria (**Fig. 1** and **Supplementary table S1**).

**Fig. 1:**
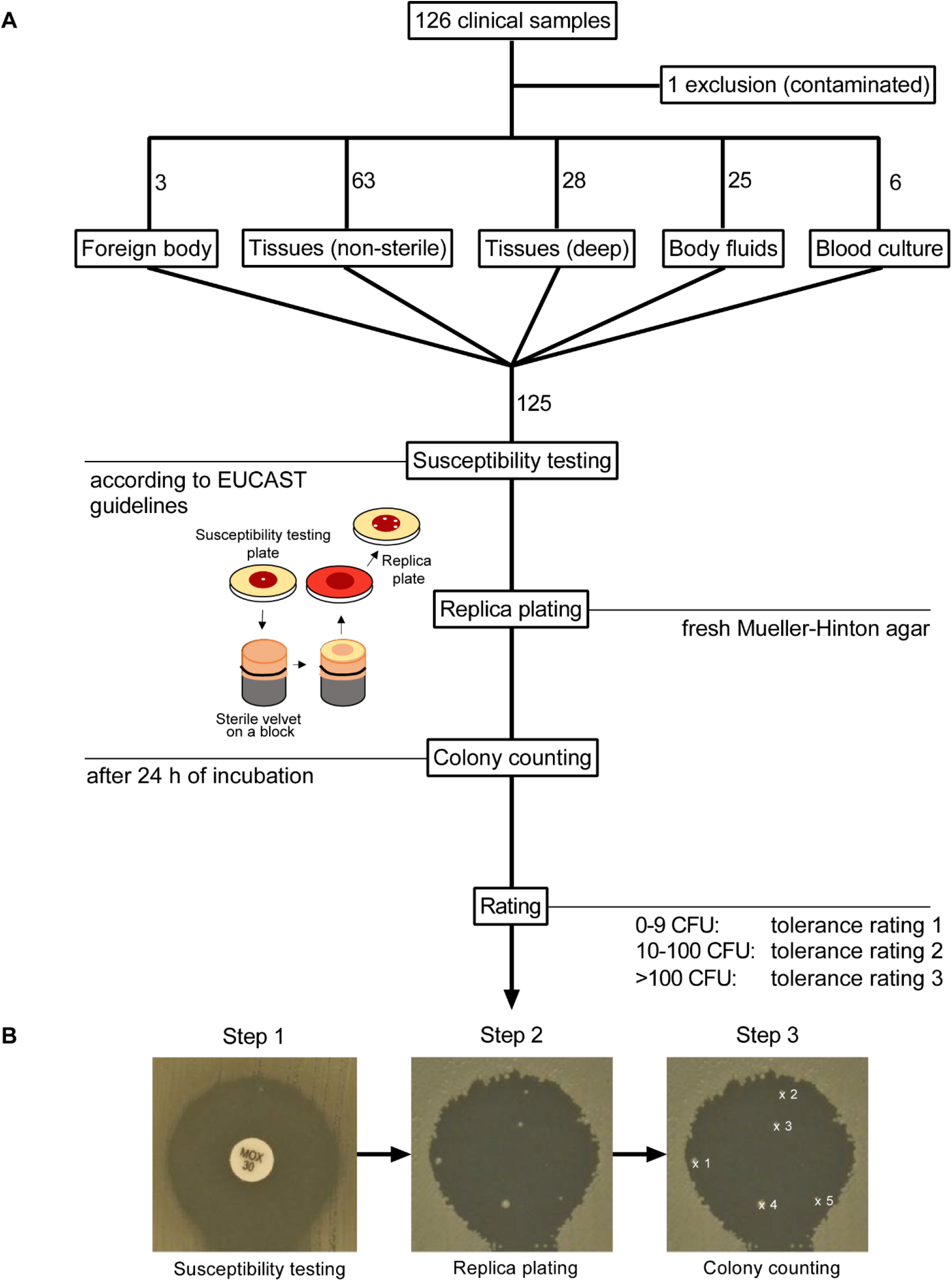
**A**) Flow chart showing the experimental procedures for all samples. The samples were divided into 5 groups based on the sampling site. Susceptibility testing was performed according to the guidelines of the European Committee on Antimicrobial Susceptibility Testing (EUCAST). The susceptibility testing plates were processed immediately after inhibition zone diameters (IZD) measurements conducted by routine diagnostic specialists. **B)** Replica plating on fresh Mueller Hinton agar allowed surviving colonies to regrow within the inhibition zones. Pictures were taken 24 and 48 hours after replica plating.

### Routine diagnostics

MALDI-TOF mass spectrometry (Bruker Daltonics, Bremen, Germany) using the direct formic acid application method (17) provided bacterial identification. Susceptibility testing and the interpretation of inhibition zone diameters (IZD) was performed according to the “European Committee on Antimicrobial Susceptibility Testing” (EUCAST) guidelines (18). The susceptibility testing plates were also evaluated phenotypically for inducible macrolide resistance (MLS phenomenon) and for the presence of the blaZ penicillinase (19), which is produced by most *S. aureus* isolates. A sharp inhibition zone edge for penicillin was taken as an index for the presence of the blaZ penicillinase (19). For statistical analysis of the impact of blaZ penicillinase production on tolerance ratings, all MRSA isolates were excluded because of narrow or missing zone of inhibition. Information about growth conditions and sample processing is provided in the **Supplementary tables S2** and **S3**.

### Replica plating

Bacterial colonies were transferred from the routine susceptibility testing plate (Mueller Hinton agar) to a fresh plate devoid of antibiotics using a sterile velvet on a metal block, thereby allowing regrowth of surviving bacteria. The plates were incubated for 48 hours at 37°C and pictures were taken at 24 and 48 hours for colony counting and regrowth rating quantification (**Fig. 1**).

### Colony counting

After replica plating, colonies growing within the former inhibition zones were counted using the ImageJ software version 1.53 (20). Regrowth was quantified as follows. <u>Tolerance rating 1:</u> 0-9 colony forming units (CFUs) within the inhibition zone, <u>Tolerance rating 2:</u> 10-100 CFUs and <u>Tolerance rating 3:</u> >100 CFUs (bacterial lawn) (15). Colonies were counted irrespective of the original IZD interpretation. In poorly delimitable inhibition zones, we only counted regrowth within 80% of the inhibition zone radius. Visually indistinguishable colony clusters were counted as one colony. If the inhibition zone on the replica plate was non-interpretable or fully overgrown on both the original and the replica plate, the antibiotic concerned was considered resistant and “not rateable”. This category was exempted from statistical analysis. Other phenomena observed are defined and documented in the supplementary materials (**Supplementary figure SF1A-D**).

### Rifampicin resistance assessment

To check for resistance development against rifampicin, we tested 18 randomly picked isolates. After replica plating, regrowing colonies within the rifampicin inhibition zone were subcultured. For each subculture, susceptibility to rifampicin was determined by disk diffusion assay and E-test (**Supplementary figure SF2)**.

### Colony-per-mm^2^ calculation

To correct for the impact of methicillin resistance on beta-lactam IZDs, we calculated the colony-per-mm^2^ ratio (CPM) for each MRSA isolate and 6 randomly picked methicillin susceptible *S. aureus* (MSSA) isolates. The original colony count (n) was divided by the inhibition zone surface, 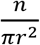.

### Clinical information

Clinical data was collected retrospectively from electronic health records (**Supplementary table S4**). For each patient, we calculated the overall regrowth score (ORS) from the replica plating assay, indicating global antibiotic tolerance. The ORS ranges from a minimum of 0 to a maximum of 39 (0 if the strain showed no regrowth at all, 39 if the regrowth rate is 3 for all 13 antibiotics). The ORS were divided into 3 groups: group 1 (low ORS: < 19), group 2 (medium ORS: 19-23) and group 3 (high ORS: ≥ 24). The groups were compared to various clinical features and the updated Charlson comorbidity index (uCCI) (21) of each patient. All samples were divided into 5 groups: blood cultures, isolates from foreign bodies (e.g. intravenous catheters), deep tissue swabs (normally sterile sites), superficial tissue swabs (sites with commensal flora), and body fluids. Details are listed in **Supplementary table S1**.

### Statistical analysis

GraphPad prism version 8.4.2. was used for statistical analysis and visualization. Spearman’s ρ, one sample t-test, Wilcoxon test, Mann-Whitney test, Fisher’s exact test and Chi-square test were used where appropriate. *p< 0.05, **p< 0.01, ***p< 0.001, and ****p< 0.0001. Statistical significance was calculated without correcting for multiple testing in exploratory analyses.

## Results

### Replica plating shows high bacterial regrowth mainly for translation targeting antibiotics and rifampicin

The replica plating of 125 clinical isolates revealed a trend towards high regrowth for antibiotics targeting protein synthesis (**Fig. 2, Supplementary tables S5 and S6**). For erythromycin, clindamycin, tetracycline and minocycline tolerance rating 2 or 3, i.e. 10 to >100 CFUs, occurred in 85.80% of all samples. For other antibiotics such as teicoplanin, tobramycin, penicillin, gentamicin, moxalactam, cefoxitin, ampicillin and norfloxacin, we observed a tendency towards low regrowth with 56.8% of all samples being rated 1. An exception was rifampicin that showed high regrowth ratings (tolerance rating 2 or 3 in 63.20%). Re-exposure of regrowing colonies to rifampicin showed no rifampicin resistance development. Results are listed in **Supplementary table S7**.

**Fig. 2:**
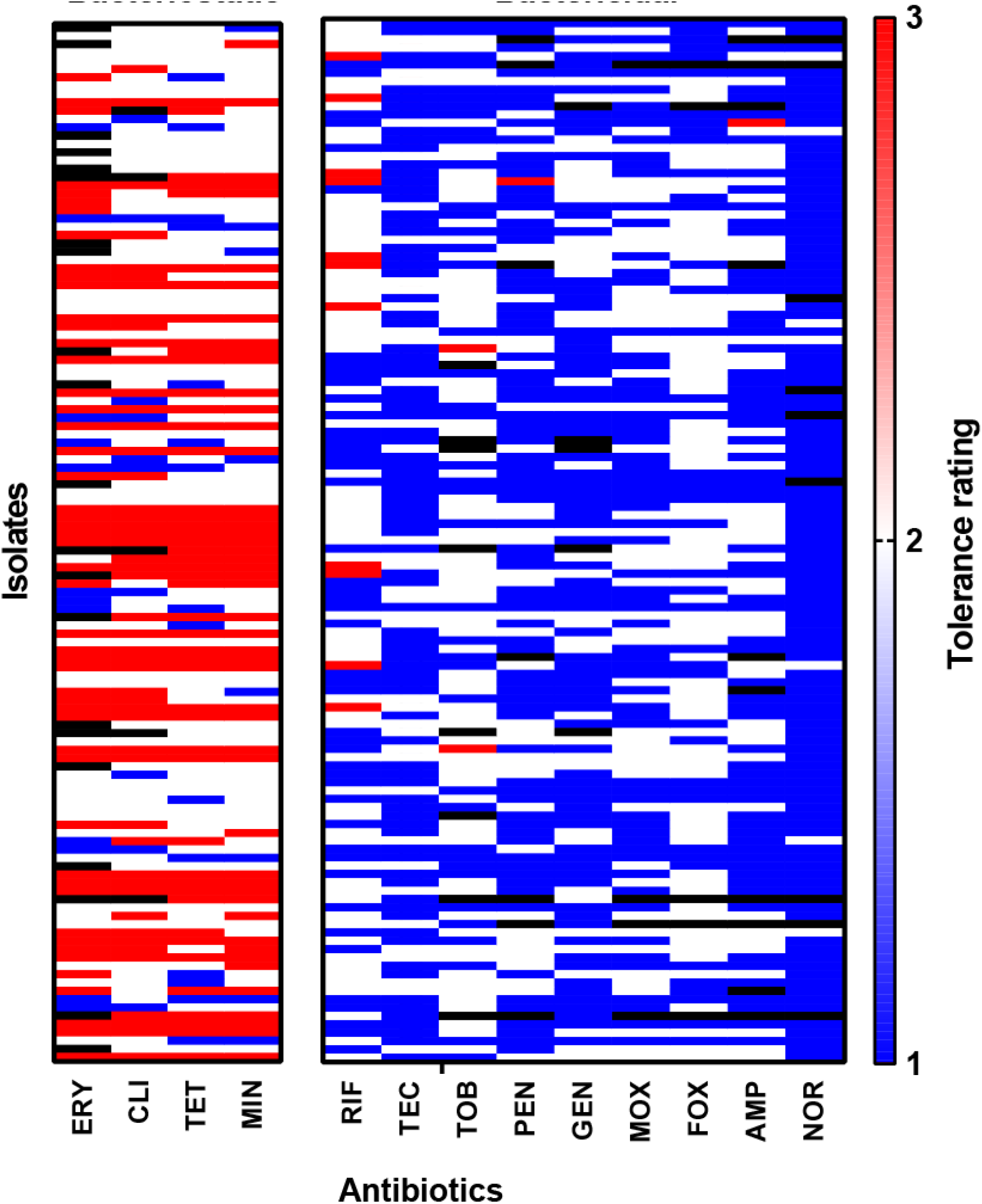
Heat map showing the regrowth ratings (1–3) of all isolates. **Antibiotics:** erythromycin (**ERY**), clindamycin (**CLI**), teicoplanin (**TEC**), tobramycin (**TOB**), penicillin (**PEN**), gentamicin (**GEN**), moxalactam (**MOX**), cefoxitin (**FOX**), tetracyclin (**TET**), minocyclin (**MIN**), ampicillin (**AMP**), norfloxacin (**NOR**), rifampicin (**RIF**). Regrowth was assessed and categorized as tolerance rating 1 (0-9 colonies within inhibition zone), tolerance rating 2 (10-100 colonies within inhibition zone) and tolerance rating 3 (>100 colonies within inhibition zone, bacterial lawn). The ratings were illustrated as a colour gradient. Not rateable isolates are marked as black boxes.

In addition to inhibition zone diameters, we investigated other phenotypical traits such as penicillinase production of the strains, as they can have an impact on the choice of antibiotic treatment. Most *S. aureus* strains produce the Ambler class A penicillinase (BlaZ) capable of hydrolysing penicillins (22), which can be phenotypically detected according to EUCAST guidelines (19).

After exclusion of all MRSA isolates, we screened 73 BlaZ-positive isolates and 45 BlaZ-negative isolates for regrowth ratings in the presence of penicillin, ampicillin, moxalactam and cefoxitin (for one isolate, information regarding BlaZ-production was not available). **Figure 3A** shows a trend for BlaZ-positive *S. aureus* towards low tolerance ratings when exposed to ampicillin and penicillin in comparison to BlaZ-negative *S. aureus*. 82.19% of all BlaZ-producing isolates (**Fig. 3B**) had a rating of one for penicillin. The same trend was observed for ampicillin, where 69.86 % of all BlaZ-producing isolates had a tolerance rating of one. Other betalactams that are not cleaved by BlaZ betalactamases did not show a similar difference: moxalactam and cefoxitin had 56.16% and 28.77% of all BlaZ-producing isolates rated 1, respectively. Details are listed in **Supplementary tables S8-S9**.

**Fig. 3:**
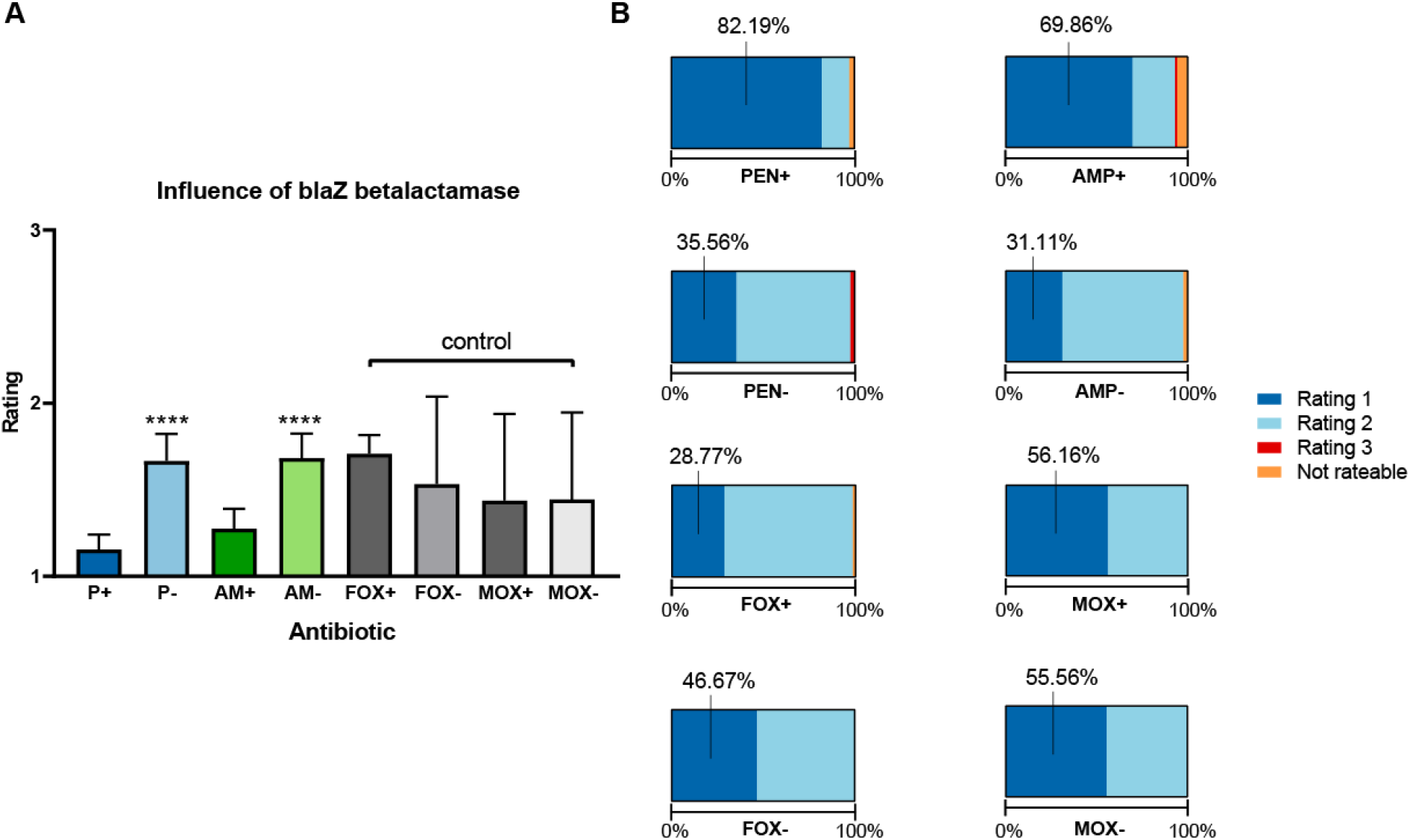
**A) Antibiotics:** penicillin (**PEN**), ampicillin (**AMP**), cefoxitin (**FOX**) and moxalactam (**MOX**). The presence (**+**) or absence (**−**) of a BlaZ penicillinase was assessed phenotypically according to EUCAST guidelines. Significance level is indicated with (*). Penicillin *p*=<0.001, ampicillin *p*=<0.001. Cefoxitin and moxalactam are not cleaved by BlaZ betalactamases and therefore serve as a control. The bars indicate the mean ratings and their 95% CI. **B)** Percentages of each rating for all betalactam antibiotics.

### Correlation analysis indicates similar effect of bacteriostatic antibiotics

Additionally, we tested for a correlation among the regrowth/ tolerance ratings for the same antibiotic class (e.g. penicillins) or among bacteriostatic antibiotics (**Table 1**). Multiple testing was not performed, as these analyses were purely exploratory. Spearman’s *ρ* showed significant correlations within all bacteriostatic antibiotics (erythromycin, clindamycin, tetracycline, minocycline). Penicillins (penicillin and ampicillin) significantly correlated with each other (*p*=<0.01, CI 95%: 0.32 – 0.62) and with moxalactam. However, there was no correlation between cefoxitin and penicillin (*p*=0.16). Furthermore, we found no correlation among aminogylcosides (gentamicin, tobramycin) or among the remaining antibiotics (further correlations are listed in **Supplementary table S10** and **S11**.

**Table 1:** Correlation matrix of tolerance ratings of tested antibiotics^a^

| Bacteriostatic antibiotics |  |  |  |  |  |  |  |  |  |
| --- | --- | --- | --- | --- | --- | --- | --- | --- | --- |
| *p*-values | | | | | Spearman's $\rho$ | | | | |
|  | ERY | CLI | TET | MIN |  | ERY | CLI | TET | MIN |
| ERY |  | <0.01 | <0.01 | <0.01 |  | 1 | 0.78 | 0.7 | 0.65 |
| CLI | <0.01 |  | <0.01 | <0.01 |  | 0.78 | 1 | 0.71 | 0.66 |
| TET | <0.01 | <0.01 |  | <0.01 |  | 0.7 | 0.71 | 1 | 0.82 |
| MIN | <0.01 | <0.01 | <0.01 |  |  | 0.65 | 0.66 | 0.82 | 1 |

| Betalactams |  |  |  |  |  |  |  |  |  |
| --- | --- | --- | --- | --- | --- | --- | --- | --- | --- |
| *p*-values | | | | | Spearman's $\rho$ | | | | |
|  | PEN | AMP | MOX | FOX |  | PEN | AMP | MOX | FOX |
| PEN |  | <0.01 | <0.01 | 0.16 |  | 1 | 0.48 | 0.36 | 0.13 |
| AMP | <0.01 |  | <0.01 | 0.03 |  | 0.48 | 1 | 0.3 | 0.2 |
| MOX | <0.01 | <0.01 |  | <0.01 |  | 0.36 | 0.3 | 1 | 0.31 |
| FOX | 0.16 | 0.03 | <0.01 |  |  | 0.13 | 0.2 | 0.31 | 1 |

| Aminoglycosides |  |  |  |  |  |  |  |  |  |
| --- | --- | --- | --- | --- | --- | --- | --- | --- | --- |
| *p*-values | | | | | Spearman's $\rho$ | | | | |
|  |  | GEN | TOB |  |  | GEN | TOB |  |  |
| GEN |  |  | 0.12 |  |  | 1 | 0.15 |  | GEN |
| TOB |  | 0.12 |  |  |  | 0.15 | 1 |  | TOB |

| Other antibiotics |  |  |  |  |  |  |  |  |  |
| --- | --- | --- | --- | --- | --- | --- | --- | --- | --- |
| *p*-values | | | | | Spearman's $\rho$ | | | | |
|  | TEC | NOR | RIF |  |  | TEC | NOR | RIF |  |
| TEC |  | 0.85 | 0.3 |  |  | 1 | -0.02 | 0.09 | TEC |
| NOR | 0.85 |  | 0.09 |  |  | -0.02 | 1 | 0.16 | NOR |
| RIF | 0.3 | 0.09 |  |  |  | 0.09 | 0.16 | 1 | RIF |

<sup>a</sup> Shown are the correlations in a matrix among the regrowth/tolerance ratings for the same antibiotic class (e.g. penicillins), or among bacteriostatic antibiotics. Multiple testing was not performed, as these analyses were purely exploratory. Significance levels are illustrated as a color gradient. Correlation was calculated using Spearman's $\rho$ . Antibiotics: erythromycin (**ERY**), clindamycin (**CLI**), tetracyclin (**TET**), minocyclin (**MIN**), teicoplanin (**TEC**), tobramycin (**TOB**), penicillin (**PEN**), gentamicin (**GEN**), moxalactam (**MOX**), cefoxitin (**FOX**), ampicillin (**AMP**), norfloxacin (**NOR**), rifampicin (**RIF**). CI: 95%.

### Clinical data

**Table 2** shows the clinical features of the patients in our cohort. We observed differences in the proportion of isolates coming from patients with immunosuppression among the overall regrowth score groups (*p*=0.0006, Chi-square test) (**Supplementary table 12**) and in the number of patients with an uCCI ≥2 (**Supplementary table 13**) (*p*=0.0309, comparing ORS group 2 and 3, Fisher’s exact test). All MRSA isolates (n=6) were found in the low overall regrowth score group (*p*=<0.0001, Chi-square test). The impact of methicillin resistance on beta-lactam IZDs was taken into consideration by calculating the CPM for each MRSA isolate and comparing it to 6 randomly selected MSSA isolates. The results are listed in **Supplementary tables S14-S19**.

**Table 2.** Baseline characteristics of patients with *Staphylococcus aureus* infections^a^

| Patient Characteristics | No. (%) of patients with characteristic |  |  |  |  |  |  |  |
| --- | --- | --- | --- | --- | --- | --- | --- | --- |
|  | Overall score <19 |  | Overall score 19-23 |  | Overall score ≥24 |  | All |  |
|  | 30 | 24% | 55 | 44% | 40 | 32% | 125 | 100% |
| Female Sex | 10 | 33.33% | 29 | 52.73% | 18 | 45.00% | 57 | 45.60% |
| Age (mean ± SD) | 31.13 | 16.71 | 48.05 | 17.17 | 71.1 | 14.8 | 50.75 | 20.87 |
| Hospitalisation | 17 | 56.67% | 18 | 32.73% | 17 | 42.50% | 52 | 41.60% |
| ICU Admission prior to isolation | 3 | 10.00% | 3 | 5.45% | 3 | 7.50% | 9 | 7.20% |
| Antibiotics 7 days prior to isolation | 13 | 43.33% | 17 | 30.91% | 13 | 32.50% | 43 | 34.40% |
| Diabetes | 8 | 26.67% | 13 | 23.64% | 12 | 30.00% | 33 | 26.40% |
| CrCl (median, IQR) | 91.5 | 56.3-101.5 | 86 | 54.5-111 | 85.5 | 51.8-111.5 | 86 | 55-109 |
| Immunosuppression | 16 | 53.33% | 10 | 18.18% | 7 | 17.50% | 33 | 26.40% |
| Malignancy | 6 | 20.00% | 7 | 12.73% | 5 | 12.50% | 18 | 14.40% |
| Foreign Material at Sampling Site | 7 | 23.33% | 7 | 12.73% | 9 | 22.50% | 23 | 18.40% |
| uCCI ≥2 | 8 | 26.67% | 14 | 25.45% | 19 | 47.50% | 41 | 32.80% |
| MRSA | 6 | 20.00% | 0 | 0.00% | 0 | 0.00% | 6 | 4.80% |
| Antibiotics 7 days prior to isolation | 7 days prior to date of isolation including systemic antibiotics and topical antibiotics at site of isolation |  |  |  |  |  |  |  |
| Diabetes | All types with or without organ-damage |  |  |  |  |  |  |  |
| CrCl | eGFR using the CKD-EPI formula (Chronic Kidney Disease Epidemiology Collaboration 2009) ml/min |  |  |  |  |  |  |  |
| Immunosuppression | Neutropenia, chronic systemic steroid use (equivalent to 20mg prednisone for 14 days) and topical steroid use at site of isolation |  |  |  |  |  |  |  |
| Malignancy | Any malignancy with or without metastasis including leukemia, lymphoma and other hematologic malignancies |  |  |  |  |  |  |  |
| uCCI | Calculated according to Ternavasio de la Vega *et al.* (18). |  |  |  |  |  |  |  |

<sup>a</sup> overview of the patient's clinical features and their uCCI in relation to the overall regrowth score (ORS); interquartile range (IQR); group 1 (low regrowth rate = ORS: < 19), group 2 (medium regrowth rate; ORS: 19-23) and group 3 (high regrowth rate; ORS: ≥ 24);

## Discussion

In this study, we analysed 125 *S. aureus* isolates derived directly from patients with various clinical presentations using the Replica Plating Tolerance Isolation System (REPTIS) and showed that it detects antibiotic tolerance in the routine diagnostic setting. In addition, REPTIS allowed identifying factors such as antibiotic regimens as well as bacterial and host features influencing the persister rate of *S. aureus*.

We found that most antibiotics targeting protein synthesis showed high regrowth ratings. In contrast, antibiotics targeting other cellular functions such as DNA-replication, transcription and cell wall synthesis, also known as bactericidal antibiotics, resulted in low regrowth rates indicating that the tested *S. aureus* isolates displayed low antibiotic tolerance levels. The results are in line with previous publications, in which bacteriostatic antibiotics exhibited steady regrowth, while bactericidal antibiotics generally had lower regrowth ratings (8). This also implies high regrowth ratings for antibiotics targeting protein synthesis, which is in line with our results. One exception was rifampicin showing high regrowth ratings of above 1 in 63.2% of all cases, making it the highest-ranking antibiotic that does not directly inhibit protein synthesis overall. Rifampicin has been used to induce persister formation probably by arresting transcription in both *E. coli* and *S. aureus* (23, 24). While being successful in inducing persister formation in *E. coli*, researchers working with *S. aureus* suspected resistance development to be the cause of regrowth (24). However, we found all isolates in this study to be rifampicin susceptible according to their inhibition zone diameter prior to replica plating. Additionally, we observed no development of rifampicin resistance in the subset of clinical isolates tested.

In contrast to rifampicin, norfloxacin exhibited strikingly low tolerance ratings (87.20% rated 1). When it was initially introduced, this quinolone was described to have less bactericidal activity against *S. aureus* as compared to ciprofloxacin (25). When measuring the MBC for *S. aureus*, other authors noted that norfloxacin had remarkably low MBC/MIC ratios (25, 26). Our results are in line with these observations.

From the bacterial side, we found that the presence of a BlaZ-betalactamase appeared to have a tendency to reduce regrowth ratings. Similarly, all MRSA isolates were in the group with low overall regrowth scores. We hypothesize that the low regrowth ratings in BlaZ-producing strains are due to the reduction of antibiotic stress on the total population. Degradation of the antibiotic compound might lead to a lower activation of stress responses and therefore reduced formation of persisters. This trend was observed for penicillin and ampicillin, both can be hydrolysed by the BlaZ penicillinase (22). However, antibiotics that are not hydrolysed as efficiently by BlaZ (e.g. cefoxitin and moxalactam) showed more ambiguous persister ratings.

Similar to these findings, we observed that all MRSA isolates were in ORS group 1 with low overall regrowth ratings. The expression of altered penicillin binding proteins (PBPs) might also reduce stress and thus explain the lower regrowth rates. However, low regrowth rates of the MRSA strains might also be caused by smaller inhibition zones leaving less space for regrowth on the replica plate. We calculated the colony-per-mm^2^ ratio of all MRSA isolates for each antibiotic and compared it to six randomly picked MSSA isolates (calculations are listed in supplementary materials, **Supplementary tables S14- S19**). Their ratios differed significantly (*p*=<0.0001, two-way ANOVA), supporting the hypothesis that altered PBPs have an impact on regrowth.

In order to assess host factors, we used the Charlson Comorbidity index (CCI) as a surrogate marker for disease severity. The CCI has recently been updated and evaluated to predict 30-day case mortality in patients with *S. aureus* bacteremia (21, 27). A cut-off value of ≥4 was found to be the best predictor for 30-day case mortality in patients with *S. aureus* bacteremia (21). Not all our patients suffered from systemic *S. aureus* bacteremia but also form localized infections which were overall less severe. We thus adjusted the uCCI cut-off value to ≥2 for our statistical analysis. Therefore, firm conclusions should be made with caution. However, it is remarkable that patients with high overall regrowth scores (≥24) more often had an uCCI of ≥2 and thus more comorbidities when compared to patients with lower overall regrowth scores. When comparing patients with low overall regrowth scores to those with medium or high overall regrowth scores, there were more immunocompromised patients in the group with low overall regrowth scores. One reason for this trend could be the persister-killing effect of cytostatic drugs that has been observed by other authors (28, 29). Two patients in our study received cisplatin as part of their treatment and the *S. aureus* isolated from these two patients had low regrowth scores. For a more detailed analysis, predefined patient cohorts will have to be tested.

Our study has several limitations. The samples were randomly included irrespective of the clinical condition. From a microbiological perspective, it makes sense to have a wide array of samples available. However, with respect to the clinical data, the statistical power may become insufficient to draw firm conclusions. To examine the clinical impact of antibiotic tolerance more accurately, predefined cohorts and samples need to be tested. Despite the low number of patients, the comparison of the results of the replica plating procedure with the uCCI showed that the former might serve as a predictor for clinical outcome in the future. The replica plating has proven to be a reliable, fast and cost-effective way of assessing *S. aureus* antibiotic tolerance in clinical microbiology laboratories and could be implicated in the future for better-tailored diagnostics.

## Acknowledgments

The authors would like to express their gratitude to the Institute of Medical Microbiology’s laboratory staff for their support in accessing the bacterial clinical isolates.

## Authors’ contributions

SCH: Experimental design, acquisition, analysis and interpretation of data, writing of the manuscript;

MH: Conceptualization, experimental design, writing and critical reading of the manuscript;

CTA: Analysis and interpretation of data, writing and critical reading of the manuscript;

FA: Critical reading of the manuscript;

AGM: Conceptualization, experimental design and critical reading of the manuscript; SMS: Critical reading of the manuscript, conceptualization;

BH: Critical reading of the manuscript, conceptualization, funding; RZ: Experimental design and critical reading of the manuscript;

SDB: Conceptualization, interpretation of data, funding, writing of the manuscript;

ASZ: Conceptualization, experimental design, data analysis, writing and critical reading of the manuscript, funding;

All authors revised the manuscript and approved the final version.

